# Inhibition of *Clostridioides difficile-*specific DNA adenine methyltransferase CamA by analogs of *S*-adenosyl-L-methionine

**DOI:** 10.1101/2025.08.13.670177

**Authors:** Jujun Zhou, Youchao Deng, Dan Yu, Taraneh Hajian, Masoud Vedadi, Xing Zhang, Robert M. Blumenthal, Rong Huang, Xiaodong Cheng

**Affiliations:** Department of Epigenetics and Molecular Carcinogenesis, University of Texas MD Anderson Cancer Center, Houston, TX 77030, USA; Borch Department of Medicinal Chemistry and Molecular Pharmacology, Purdue Institute for Drug Discovery, Purdue Institute for Cancer Research, Purdue University, West Lafayette, IN 47907, USA; Drug Discovery Program, Ontario Institute for Cancer Research, Toronto, Ontario M5G 0A3, Canada; Department of Pharmacology and Toxicology, University of Toronto, Toronto, Ontario M5S 1A8, Canada; Department of Medical Microbiology and Immunology, and Program in Bioinformatics, The University of Toledo College of Medicine and Life Sciences, Toledo, OH 43614, USA

**Keywords:** DNA/RNA adenine methylation, adenosine analogs, *Clostridioides difficile* infections, CamA, cell cycle-regulated DNA methyltransferase, CcrM, MettL5

## Abstract

Epigenetically-targeted therapies, especially those inhibiting *S*-adenosyl-L-methionine (SAM)- dependent methylations of DNA, mRNA and histones, have advanced rapidly in cancer treatment. However, these therapies remain underexplored for antibiotic development, despite the growing threat of antimicrobial resistance. Here, we screened a focused library of SAM analogs against the DNA adenine methyltransferase CamA specific to the enteric pathogen *Clostridioides difficile.* At the same time, we examined six other adenine methyltransferases, including two bacterial DNA methyltransferases, and four human RNA methyltransferases having distinct RNA substrates. Compound **113** selectively inhibited CamA (IC_50_ = 0.15 μM). In addition, compound **67** inhibited *Caulobacter crescentus* CcrM (IC_50_ = 1.8 μM), which has orthologs present in pathogens such as *Brucella*; while compounds **77** and **37** inhibited the human DNA methyltransferase complexes MettL3-MettL14 and MettL5-Trm112, respectively, at 7-8 μM concentrations. These results provide chemical probes for exploring the role of CamA in sporulation and colonization, with potential as antivirulence agents against *C. difficile* infection. Our study also introduces the first chemical probes for inhibiting bacterial CcrM and human MettL5, each of which plays key roles in their respective hosts.

**Table of Contents Graphic (TOC):** 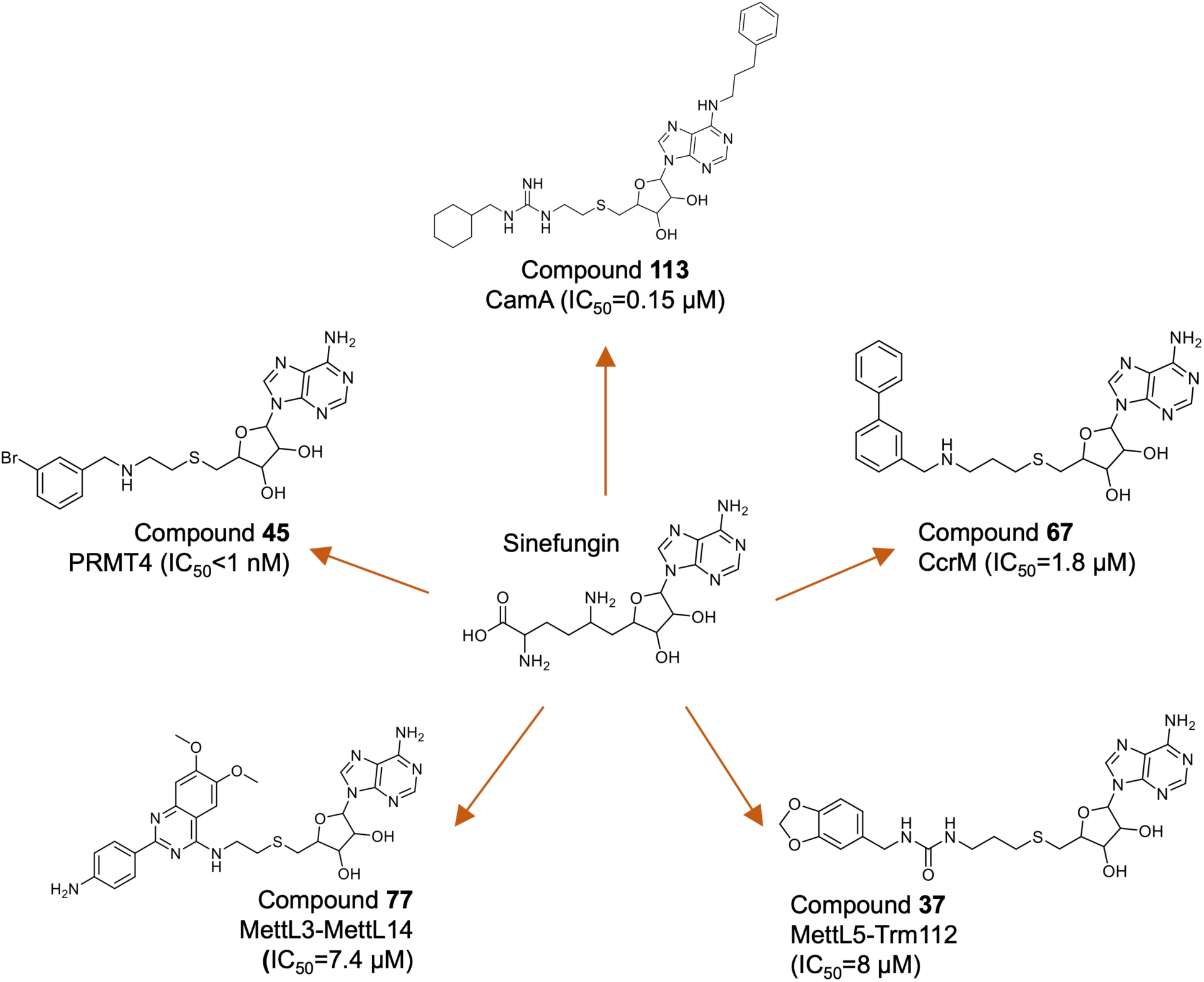

## INTRODUCTION

*Clostridioides difficile* infection (CDI) is a leading cause of community-associated infection in children, and of hospital-acquired diarrhea in older patients ^1-3^. CDI can have life-threatening consequences and is considered to be an urgent threat to public health. According to a recent report by the Centers for Disease Control and Prevention, the incidence rate of CDI increases with age, is higher in women than in men, and is more prevalent among Caucasian individuals compared to other racial groups ^4^. At present, no vaccine is available ^5^, and one of the major treatments – Fecal Microbiota Transplantation (FMT) – is still in the process of being characterized and optimized ^6,7^. Phage therapy and monoclonal antibody treatments are also under development ^8^. In addition, several small-molecule therapeutics targeting *C. difficile* have progressed to phase I-III clinical trials ^9^. These include CRS3123 (a tRNA^Met^ synthetase inhibitor) ^10^, ibezapolstat (a DNA polymerase IIIC inhibitor) ^11, 12^, and two DNA <u>m</u>inor <u>g</u>roove <u>b</u>inders - MGB-BP-3 ^13^ and ridinilazole ^14^. Thus, while several potentially-beneficial therapeutic approaches are under active development, the fact remains that there is no consistently-efficacious *C. difficile* treatment at present. Furthermore, antivirulence drugs – which target the pathogenesis rather than the growth of a bacterium – are less likely to select for antibiotic resistance ^15^, and can synergize with more traditional antibiotics ^16^.

Genomic analyses of sequenced *C. difficile* strains have identified a DNA adenine methyltransferase (CamA) unique to, and widespread within, this species ^17, 18^. CamA is essential for normal sporulation, and is required for persistent CDI in an animal model, but not for *C. difficile* growth ^17^. Because recurrent CDI is often associated with antibiotic use within the prior 12 weeks ^4, 19^, and because *C. difficile* itself is increasingly antibiotic resistant ^20, 21^, there is a pressing need for non-conventional therapeutic agents to effectively control CDI, and CamA appears to be a potentially valuable target.

CamA is a member of the *S*-adenosyl-*L*-methionine (SAM)-dependent methyltransferase (MTase) family, which catalyzes methyl transfers to a broad range of substrates ^22^, including DNA, RNA, proteins, and small molecules such as histamine and thiopurine ^22^. SAM is a metabolite synthesized in nearly all species ^23, 24^, with the exceptions being obligate parasites that obtain SAM from their hosts ^25, 26^. SAM is by far the major methyl donor in transmethylation reactions ^27, 28^, and also participates in radical SAM reactions ^29-33^, polyamine biosynthesis ^34^, and SAM-sensing riboswitch regulation ^35-37^. Given the importance of nucleic acid and histone methylation in diseases such as cancer and immune disorders ^38-41^, SAM-dependent MTases have become attractive therapeutic targets ^42-47^. SAM analogs are versatile tools for probing and inhibiting MTase activity ^48-53^, with some of them already in use in clinical trials or preclinical studies that target human epigenetic enzymes ^54, 55^ and viral RNA MTases ^56-59^.

We previously demonstrated that several SAM analogs, originally designed as selective inhibitors of human protein arginine MTases (PRMTs) ^55, 60, 61^ and of the histone H3 lysine 79 MTase DOT1L ^62^, also inhibit CamA *in vitro* at low micromolar concentrations ^63^. Subsequent optimization of adenosine analogs yielded CamA inhibitors with half-maximal inhibitory concentration (IC_50_) values in the sub-micromolar range ^64, 65^.

In the present study, we leveraged a focused library of SAM analogs to screen against three bacterial DNA adenine MTases: our primary target of interest, *C. difficile* CamA, which modifies the AT-rich CAAAA**<u>A</u>** sequence, *Escherichia coli* Dam, which methylates G**<u>A</u>**TC sites ^66^ and is widespread among γ-Proteobacteria ^67, 68^, and *Caulobacter crescentus* cell cycle-regulated DNA MTase (CcrM), which targets the G**<u>A</u>**nTC motif (n = any nucleotide). While *C. crescentus* is not itself a human pathogen, CcrM orthologs play key regulatory roles in other α-Proteobacteria, including pathogens like *Brucella* ^69, 70^ and the colorectal cancer-associated *Fusobacterum nucleatum* ^71^. To test for selectivity, we also screened this SAM analog library against four human RNA adenine MTases, each acting on distinct RNA substrates. Finally, we discuss the potential of small-molecule therapeutics targeting *C. difficile* in relation to CamA inhibition.

## RESULTS

### A library of SAM analogs

We began with a set of pan-PRMT inhibitors ^65, 72^ and a focused library of adenosine analogs, which led to the identification of a PRMT4-selective inhibitor with an IC_50_ < 0.5 nM ^73^, and ultimately yielded a curated library of 114 compounds (Figure 1). All compounds share a 5’-thioadenosine (S-Ad) scaffold with a flexible 2-3 carbon linker (Figure 1A-J). Compounds in groups A and B incorporate a moiety that resembles the guanidino moiety of the arginine side chain (Figure 1A-B), while groups C and D replace the guanidino with a urea moiety (Figure 1C-D). Groups E and F feature a benzylamine with a single hydrogen bond donor (NH) (Figures 1E-F). Group G compounds contain a dimethoxyquinazolin-4-amine moiety (Figure 1G), group H includes a phenyl ether moiety (Figure 1H), and group I introduces a phenylpropane moiety at the N6 position of the adenosine ring ^64^ and a 3-carbon linker (Figure 1I), whereas compound **113** contains a 2-carbon linker (Figure 1J). As a control, sinefungin (**114**) ^74^, a pan-inhibitor of SAM-dependent MTases, was included in the screen (Figure 1K), bringing the total library size to 114 compounds (Supplementary Table S1). As compounds **38** and **42** were synthesized from different batches but have the same structure, they served as an internal quality control. Thus, there are a total of 113 unique structures in this library.

**Figure 1.**
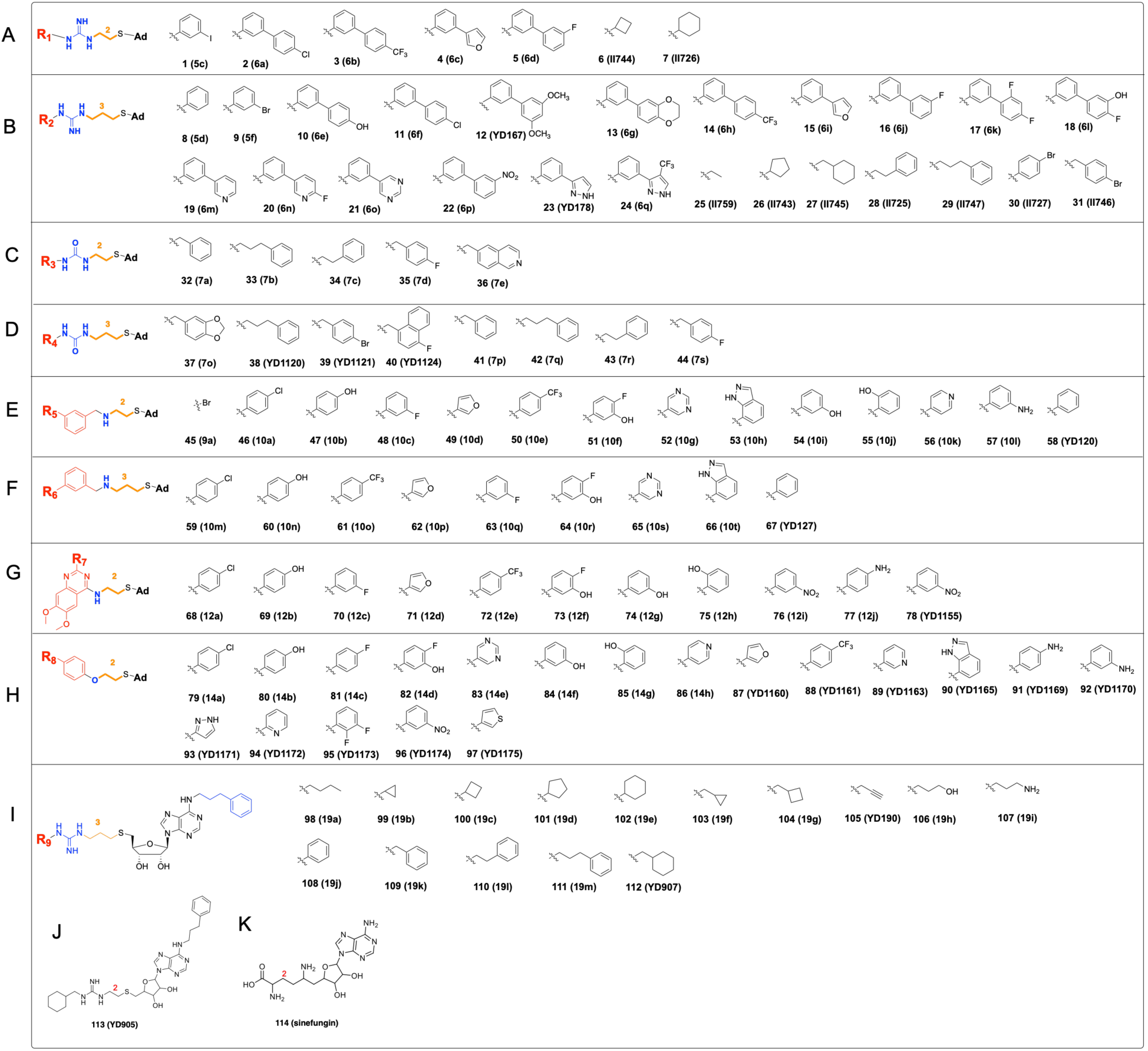
A library of 114 SAM analogs sharing a 5’-thioadenosine scaffold with a 2-3 carbon linker (see Supplementary Table S1). **(A-B)** Compounds incorporate a guanidino moiety. (**C-D**) Compounds feature a urea moiety. We note that compounds **38** and **42** are the same compound synthesized twice. (**E-F**) Compounds feature a benzylamine. (**G**) Compounds contain a dimethoxyquinazolin-4-amine moiety. (**H**) Compounds include a phenyl ether moiety. (**I**) Compounds introduce a phenylpropane moiety at the N6 position of the adenosine ring. (**J**) Compound **113**. (**K**) Sinefungin (**114**).

### Library screening against seven DNA or RNA adenine methyltransferases

We tested the SAM analog library against three bacterial DNA MTases and four human RNA MTases that act on distinct RNA substrates (Figure 2A). All seven enzymes catalyze methylation of the exocyclic amino group of adenine at the N6 position, producing N6-methyladenine. We reasoned that, during the methylation reaction, the adenine ring’s exocyclic amino group may undergo chemical transformation similar to that of the planar guanidino group of arginine that is targeted by PRMTs. An initial inhibition screen was conducted at a compound concentration of 10 μM using the MTase-Glo biochemical assay, which measures the conversion of the methyl donor SAM to *S*-adenosyl-L-homocysteine (SAH) ^75, 76^.

**Figure 2.**
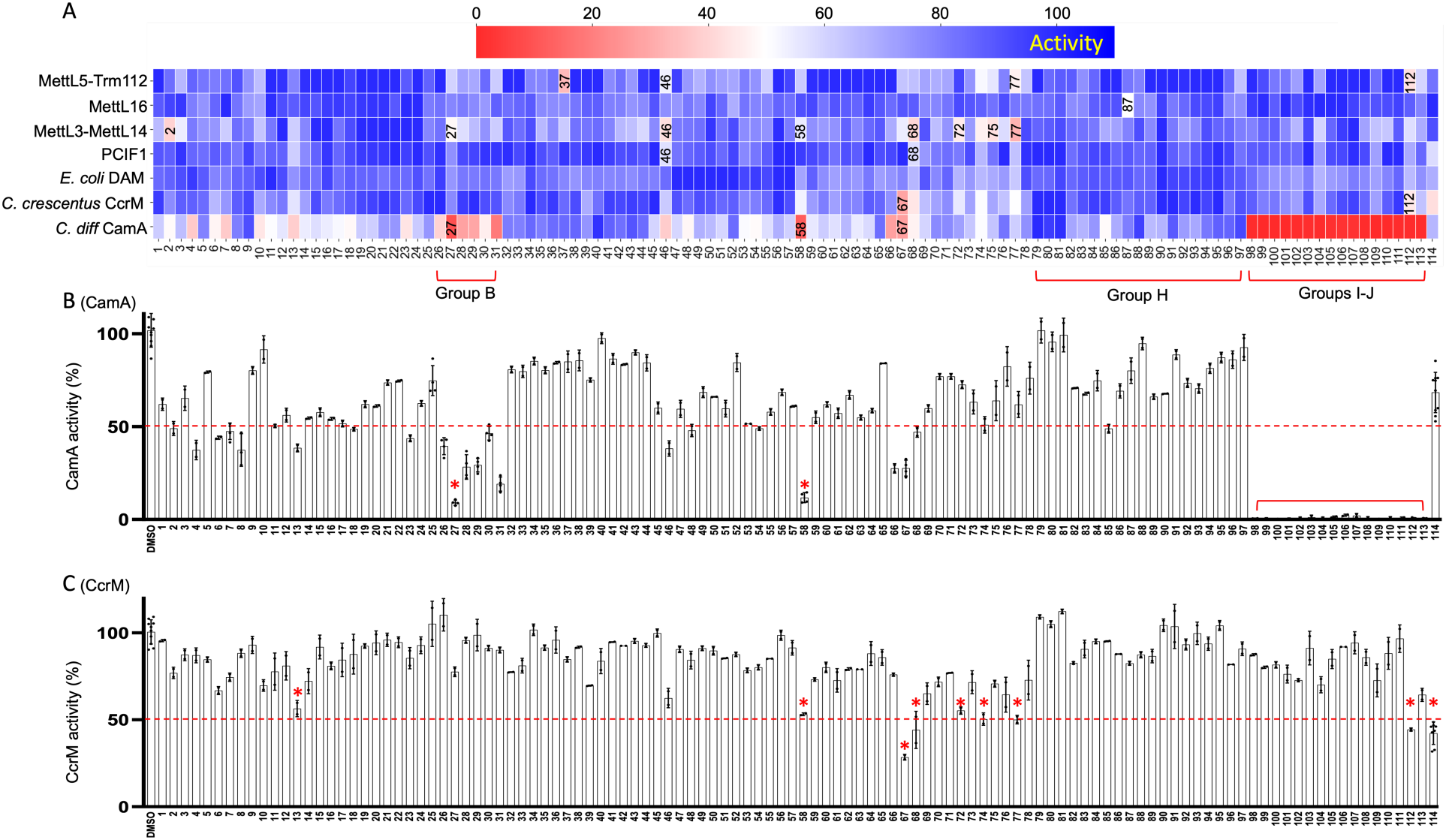
Inhibition study of seven MTase activities at a single inhibitor concentration of 10 μM. (**A**) Heatmaps of relative inhibition by compounds against four human RNA MTases and three bacterial DNA MTases, where redder shades indicate increasing inhibition. (**B**) Relative inhibition by compounds against *C. difficile* CamA in a bar graph. N = 2 replicates. (**C**) Relative inhibition by compounds against *C. crescentus* CcrM in the bar graph. N = 2 replicates. The three panels are vertically aligned by compound. For other inhibition data, see Table S1.

Among the three bacterial DNA MTases tested (Dam, CcrM and CamA), no compounds meaningfully inhibited (*i.e.*, >50%) *E. coli* Dam, which methylates the G**<u>A</u>**TC sequence. In contrast, seven compounds inhibited *C. crescentus* CcrM (which targets G**<u>A</u>**nTC motifs) by 50% or more. Surprisingly, 18 compounds inhibited CamA activity by more than 90%. Based on these initial results, *E. coli* Dam was excluded from further analysis.

### Inhibition of CamA

The 18 compounds that inhibited CamA activity by more than 90% at 10 μM are **27** and **58**, and all sixteen compounds from groups I-J (Figure 2B). Groups I-J compounds share a common structural feature: a phenylpropane moiety attached at the N6 position of the adenosine ring (Figure 1I-J). This observation suggests that analog N6-substitution, as opposed to modifications at the SAM homocysteine moiety, provides greater potency and selectivity for targeting CamA. This result aligns with prior structural insights showing that SAM binding to CamA involves a conformational rearrangement of two N-terminal helices ^77^, and that the solvent-exposed edge of the SAM adenosine moiety can be derivatized to make inhibitory analogs ^64^. Compared to parent compounds in Group B, N6-substituted analogs showed greater inhibition, indicating the important contribution from 3-phenylpropyl group for interacting with CamA. Since no compound in Group A showed any promising inhibition, it implies that a 3-C linker is preferred. Selective inhibition of CamA may also reflect the fact that CamA has an unusually low binding affinity for SAM, relative to most other MTases (*K*_m_ in the range of 17-25 μM) ^65, 77^. For comparison, *E. coli* Dam has a SAM *K*_m_ of 3-6 µM ^78-80^, while the human enzymes PCIF1 and MettL5 exhibit SAM *K*_m_ values of 0.7 µM and 1.0 µM, respectively ^81^. However, this rationale does not explain the lack of inhibition of MettL16, which has a reported SAM *K*_m_ > 400 µM ^81^. Compared to Groups A-D, a guanidine moiety appears to be preferred over a urea. There is no hit from Group H except **85,** which showed ∼50% inhibition, suggesting that substitution at the para-position of the phenyl ring is disfavored.

Next, we selected four SAM analogs (compounds **27**, **58**, **67** and **113** in Figure 3A) for more-detailed inhibition studies, along with the previously-characterized adenosine analog MC4741 for comparison. MC4741 lacks the homocysteine moiety but, like compound **113**, carries a 3-phenylpropyl group at the N6-amino position of adenosine. Compound **67** was also chosen for its overlapping activity against CcrM. Under the defined conditions with 0.05 μM CamA, 40 μM SAM, and 2.5 μM DNA substrates, the five compounds exhibited IC_50_ values ranging from 9 - 0.15 µM (Figure 3B). Reducing the linker length from three carbons (**67**) to two carbons (**58**) more than doubled the potency. Substituting the secondary amine in compounds **67** and **58** with a guanidino-containing moiety (compound **27**) further increased potency 2.7X. MC4741, despite lacking the homocysteine moiety (Figure 3A), showed twice the potency of compound **27** (Figure 3C), suggesting that the N6 substitution plays a key role. Compound **113**, which contains both the guanidino group and the 3-phenylpropyl moiety, exhibited the highest potency with an IC_50_ of 0.15 µM. Structural comparison revealed that compounds **113** (PDB 8FS1) and MC4741 (PDB 8CXZ) occupy the SAM-binding pocket in a manner similar to sinefungin (PDB 7RFK), with the adenosine ring aligned closely among these structures (RMSD ≤ 1 Å between the protein components; Figure 3D-F).

**Figure 3.**
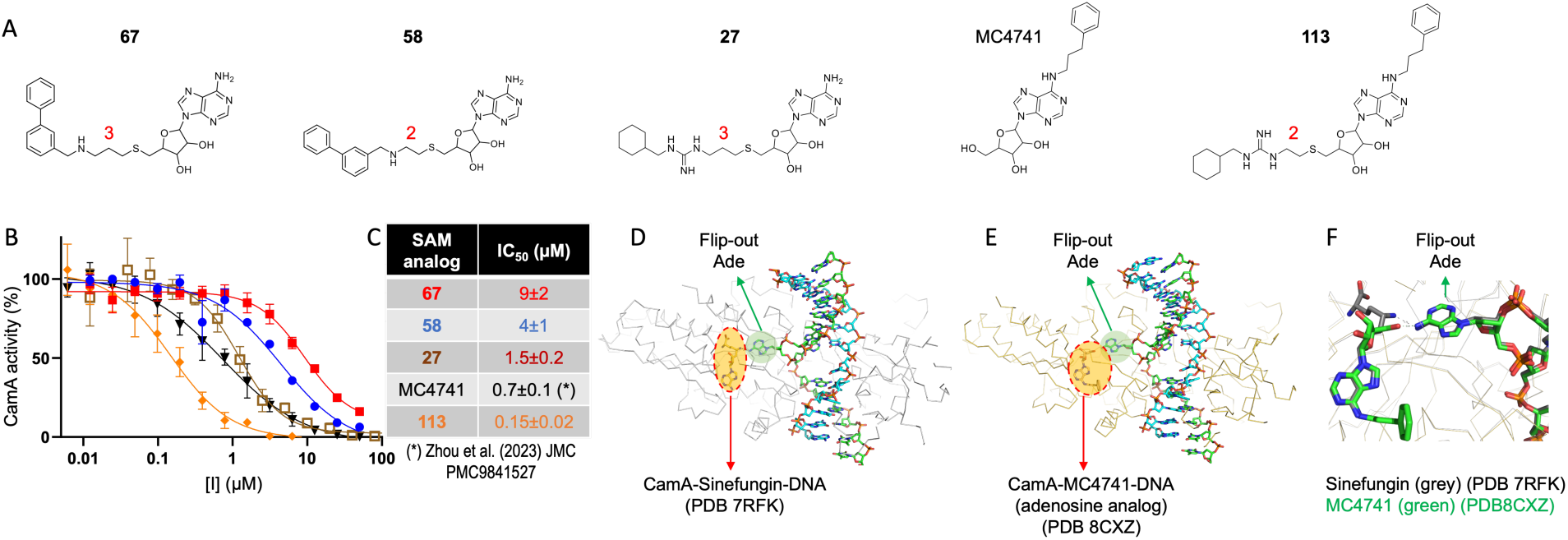
Inhibition of CamA. **(A)** Chemical structure of the five SAM analogs used most in this study. (**B**) The IC_50_ measurements as a function of inhibitor concentrations. Error bars indicate the mean ± SD of N=3 independent determinations. (**C**) Summary of the IC_50_ values. (**D**) Structure of CamA-sinefungin-DNA (PDB 7RFK). (**E**) Structure of CamA-MC4741-DNA (PDB 8CXZ). (**F**) Overlap of sinefungin and MC4741.

### Inhibition of CcrM

As shown in Figure 2C, a single-dose screen at 10 µM revealed that, in order of decreasing potency, compound **67** inhibited *C. crescentus* CcrM activity by 72%, followed by **112** (56%), **74** (50%), **77** (50%), **58** (47%), and **13** (44%). Besides compound **67**, all five others share similar potency to that of Sinefungin (**114**). The IC_50_ of compound **67** was determined to be 1.8 µM, approximately1.7X more potent than sinefungin (Figure 4A-B). Unlike CamA, which functions as a monomer, CcrM is a β-class MTase and operates as a homodimer ^82^: one subunit binds and recognizes the target DNA strand and catalyzes the methyl transfer, while the second subunit interacts with the non-target strand ^83^ (Figure 4C). Interestingly, compound **67** exhibited 5X stronger inhibition of CcrM compared to CamA, with IC_50_ values of 1.8 µM and 9 µM, respectively. Using the Protenix server ^84^, an open-source reproduction of AlphaFold3 ^85^, we modeled the binding of compound **67** to CcrM with high confidence scores of both <u>p</u>redicted template <u>m</u>odeling (pTM) and interface <u>p</u>redicted template <u>m</u>odeling (iPTM) metrics (Figure 4D). Among the top five predicted models, compound **67** consistently occupies the SAM binding pocket, showing alternative conformations of its biphenyl moiety (Figure 4D).

**Figure 4.**
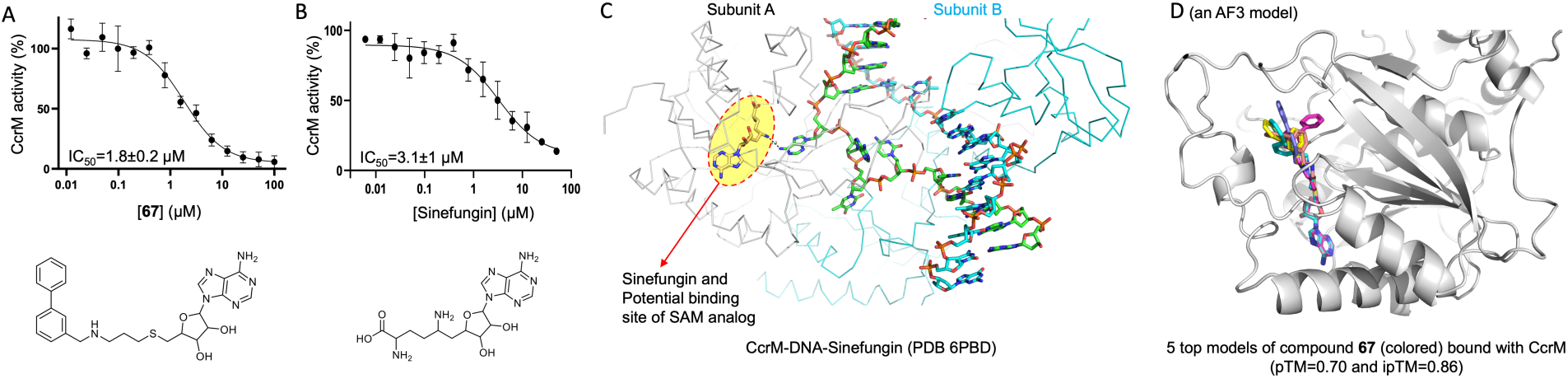
Inhibition of CcrM. (**A-B**) The IC_50_ measurements of SAM-analog **67** and sinefungin (**114**). Error bars indicate the mean ± SD of N=3 independent determinations. (**C**) Structure of CcrM-sinfungin-DNA (PDB 6PBD). (**D**) AlphaFold3 predicted models of compound **67** bound with CcrM.

### Inhibition of human RNA MTases

Among the four human RNA enzymes tested, MettL5 targets rRNA, MettL16 modifies snRNA, with MettL3 and PCIF1 methylating mRNA at different sites. Dysregulation of these RNA modifications can alter gene expression ^86^, particularly in cancer cells ^87^. MettL16 methylates adenine within a conserved UAC**<u>A</u>**GAGAA sequence, found in hairpins of the 3’ UTR of the SAM synthetase (MAT2A) mRNA and in U6 snRNA ^81, 88-90^. Among all tested compounds, only one (compound **87**) showed ∼50% inhibition of MettL16 (Figure 2A).

PCIF1 (<u>p</u>hosphorylated RNA polymerase II <u>C</u>TD interacting <u>f</u>actor <u>1</u>) ^91^ methylates adenosine when it is the first transcribed nucleotide after the mRNA cap ^92-95^. Two compounds, **46** and **68**, showed ∼40% inhibition of PCIF1 (Figure 2A) and contain a para-chlorobenzene moiety. Notably, compound **46** inhibited PCIF1 (42% inhibition), MettL5 (46%), and MettL3 (58%) to similar extents, while compound **68** also showed overlapping inhibition of PCIF1 (42% inhibition), MettL3 (60%), and MettL5 (35%) (Figure 2A).

### Inhibition of MettL5-Trm112

MettL5 forms a heterodimer with Trm112 and catalyzes adenine methylation at position 1832 of 18S rRNA ^81, 96-100^, modulating translation of mRNA ^101^. Of the 120 compounds tested, three compounds - **37** (66% inhibition), **77** (53%) and **112** (57%) - showed greater than 50% inhibition of MettL5-Trm112 at 10 μM (Figure 2A). We determined an IC_50_ value of 8 µM for compound **37** against MettL5-Trm112 activity (Figure 5A). We note that compound **37** is a selective inhibitor of MettL5-Trm112 among the six enzymes tested (Figure 2A). Given that MettL5 affects differentiation of embryonic stem cells ^97^, and promotes tumorigenesis in multiple cancer models ^87, 98, 102-105^, compound **37** may serve as a promising lead for further optimization as an anticancer therapeutic. AlphaFold3-based prediction from the Protenix server positioned compound **37** within the SAM-binding site of MettL5 (Figure 5B). Notably, four out of the five top models show a bent conformation, with their 1,3-benzodioxole moiety extending into the substrate-binding site (Figure 5C). This orientation effectively bridges between the SAM-binding and substrate-binding pockets, consistent with the intended design of a bi-substrate inhibitor.

**Figure 5.**
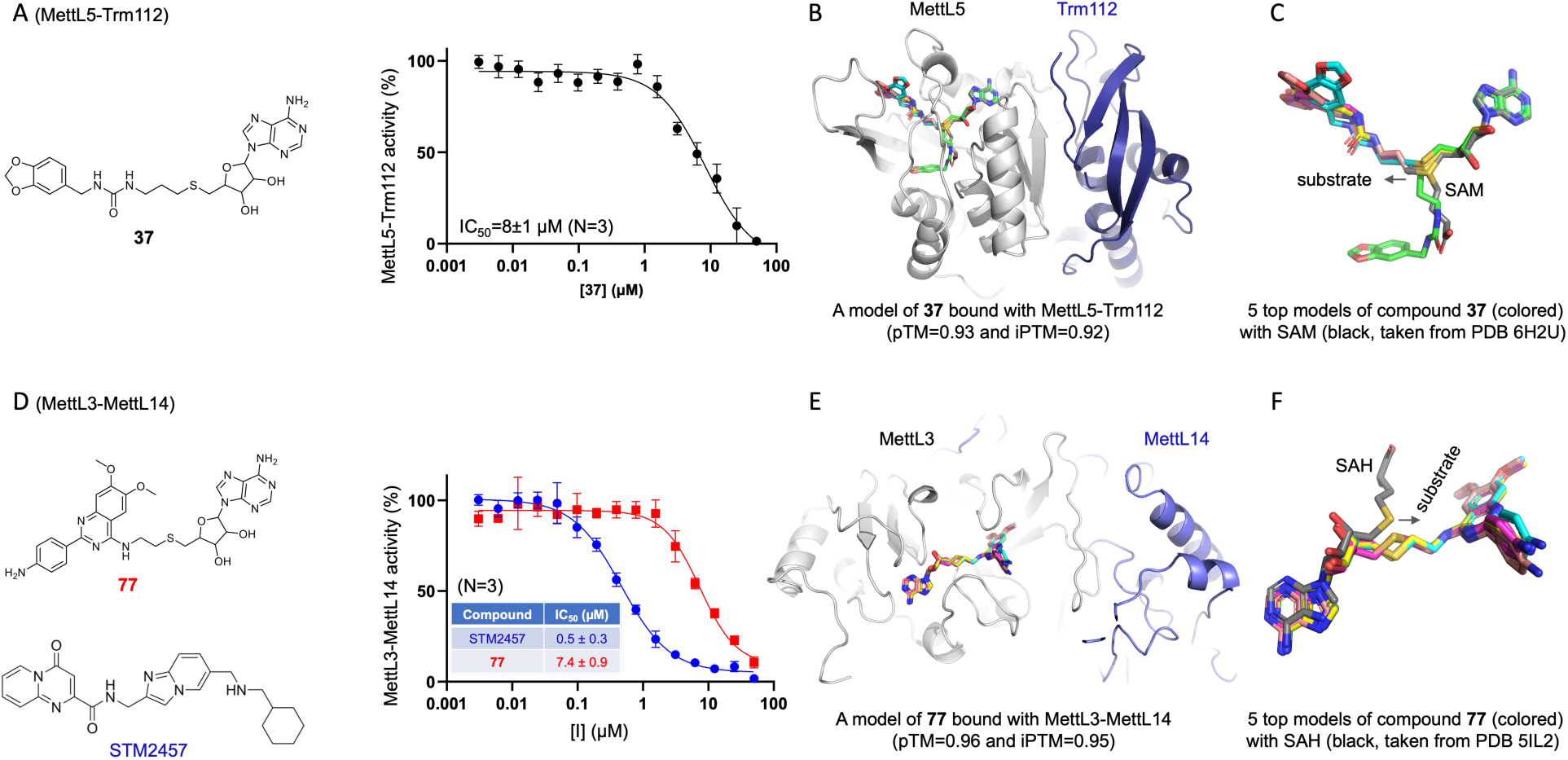
Inhibition of MettL5-Trm112 and MettL3-MettL14. (**A**) The IC_50_ measurement of compound **38** against MettL5-Trm112. Error bars indicate the mean ± SD of N=3 independent determinations. (**B**) AlphaFold3 predicted models of compound **37** bound with MettL5. (**C**) Four of the five top models adopt a bent conformation, while one model (green) aligns with the homocysteine moiety of SAM. The arrow indicates the direction of methyl transfer to the substrate. (**D**) The IC_50_ measurements of compounds **77** and STM2457 against MettL3-MettL14 were made by varying inhibitor concentrations. Error bars indicate the mean ± SD of N=3 independent determinations. (**E**) AlphaFold3 predicted models of compound **77** bound with MettL3. (**F**) Five top models bridge between the SAM-binding and substrate-binding pockets. The arrow indicates the direction of methyl transfer to the substrate.

### Inhibition of MettL3-MettL14

Like MettL5-Trm112 heterodimer, MettL3 also forms a heterodimer, pairing with MettL14 ^106, 107^, and catalyzing adenine-N6 methylation in mRNA (in this case at the degenerate consensus sequence RR**<u>A</u>**CH, where R = purine and H * G) ^108^. Also like MettL5-Trm112, MettL3-MettL14 affects embryonic stem cell differentiation ^109^ and can promote tumorigenesis ^110-113^. Unlike MettL5-Trm112, the MettL3-MettL14 complex is also active on DNA substrates, at least *in vitro* ^114, 115^. Seven SAM analogs exhibited 50-60% inhibitory activity against MettL3-MettL14 using an RNA substrate (Table 1): **2** (60% inhibition), **27** (52%), **46** (58%), **58** (59%), **68** (54%), **72** (53%), **75** (57%), while compound **77** displayed 68% inhibition. Other MettL3 inhibitors have been reported ^116^. We compared the potency of compound **77** to a known MettL3 inhibitor, STM2457 ^117^, which target the SAM-binding site. Compound **77** showed inhibitory activity at IC_50_ of 7.4 µM, which was less potent than STM2457 (Figure 5D). Nevertheless, compound **77** was modeled as a bi-substrate inhibitor occupying both SAM-binding and substrate-binding pockets (Figure 5E-F).

**Table 1.**
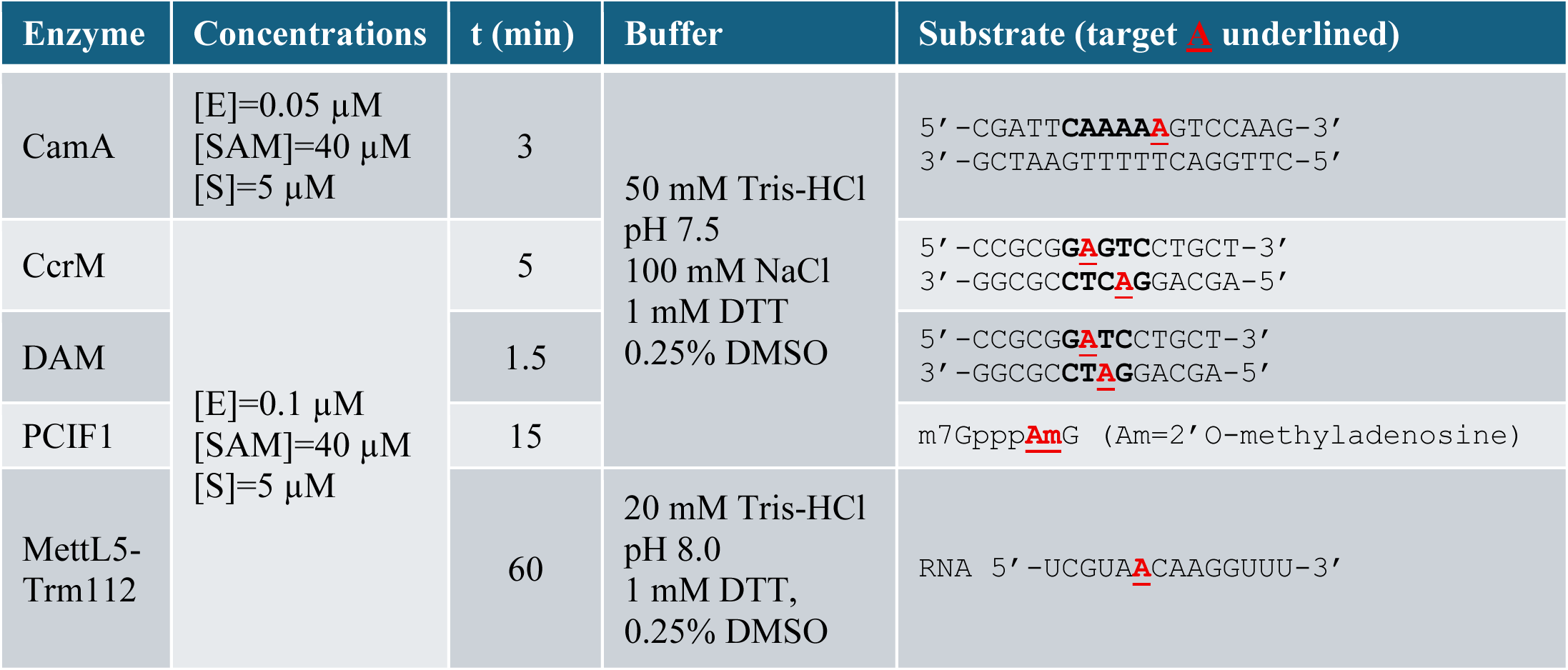
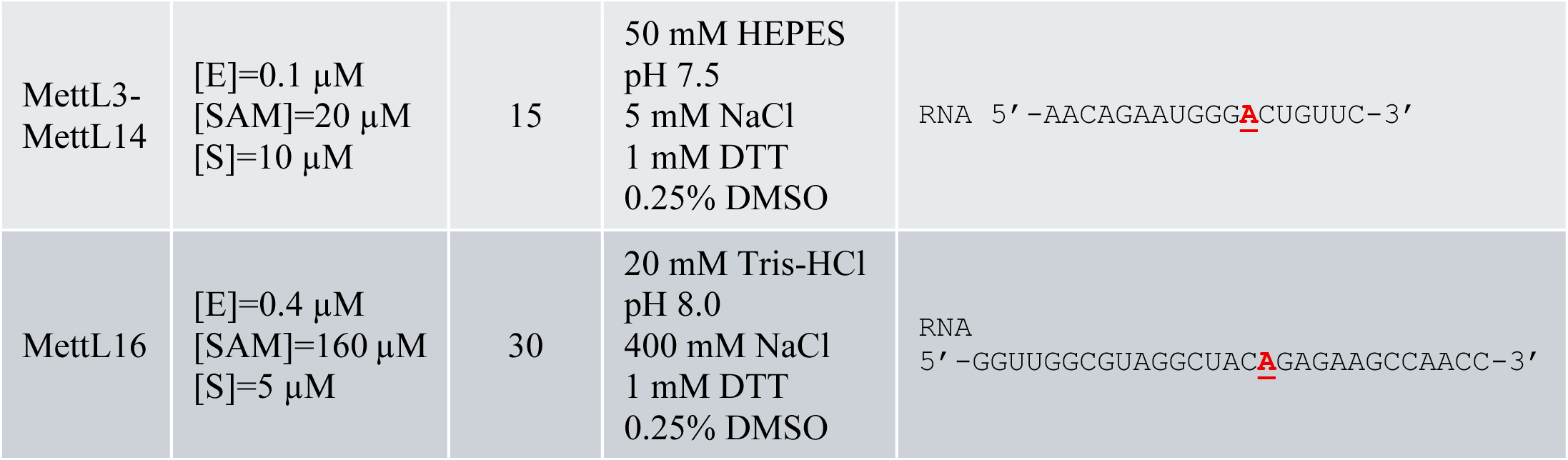
DNA or RNA methylation reaction conditions at room temperature (∼21 °C)

## DISCUSSION

### Inhibition of nucleic acid MTases

Using our focused SAM analog library, we previously identified a selective inhibitor (compound **45**) of human PRMT4/CARM1 ^73^ (Figure 6A). In the current study, we identified unique scaffolds of SAM analogs that inhibit three additional MTases *in vitro* with varying potency: compound **113** as a selective inhibitor of *C. difficile* CamA (Figures 3 and 6B), compound **67** for *C. crescentus* CcrM, and compound **37** for human MettL5 (Figure 5). These compounds represent promising leads for further optimization toward their respective targets. In the case of **113**, this could lead to development of a therapeutic agent against *C. difficile*, which can cause lethal infections and is difficult to treat ^118^. For **67**, this may lead to agents targeting the CcrM orthologs in frank pathogens such as *Brucella* ^119^ and the colorectal cancer-associated *Fusobacterum nucleatum* ^71^. For **37**, this might lead to a useful anticancer drug ^102^. Among all nine chemical groups, Group H (containing a phenylether group) is less favorable for all tested MTases, except that compounds **85** and **87** respectively showed a modest inhibition of CamA and MettL16. Groups I and J (with N6 phenylpropyl substitution) clearly, selectively, and potently inhibited CamA, indicating a unique binding pocket in CamA in the studied group. Among all tested MTases, CamA and MettL3-MettL14 are relatively more tolerant for binding these compounds, as more hits were identified, while the remaining MTases have stricter preferences. PCIF1 favors a para-chlorophenyl group as shown in both hits (compounds **46** and **68**). Compound **67** showed dual inhibition of both CcrM and CamA. Interestingly, its analog **58** (with a 2-C rather than 3-C linker) showed increased potency and selectivity to CamA.

**Figure 6.**
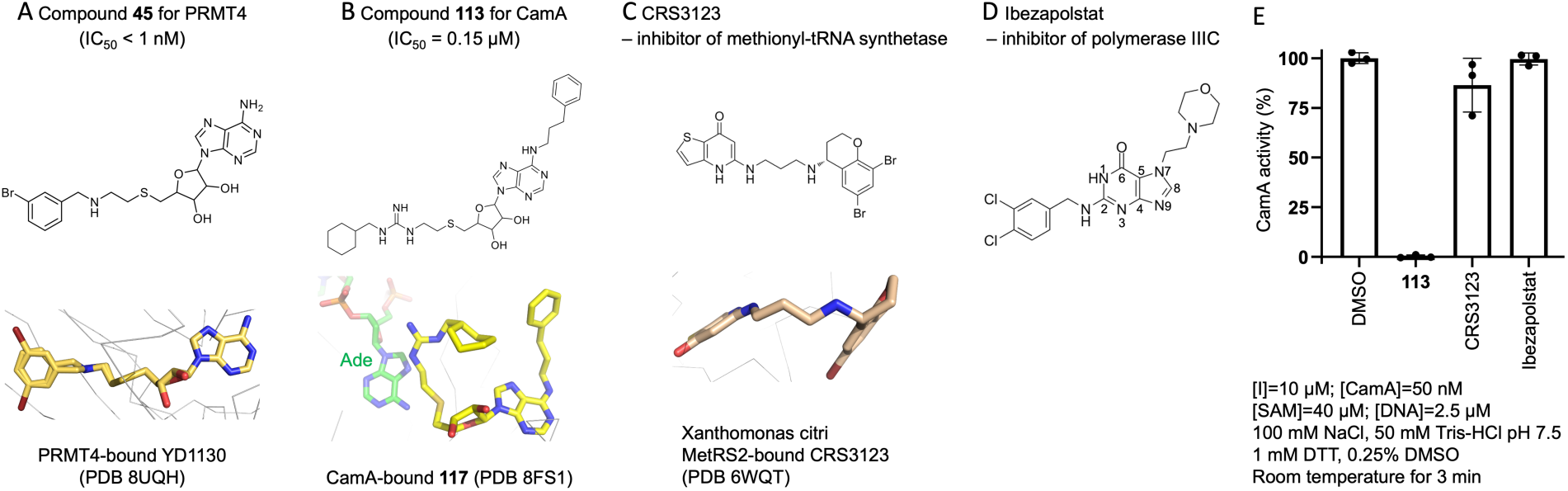
Comparison with existing antibacterial compounds. (**A**) Compound **46** for PRMT4, (**B**) compound **113** for CamA. (**C**) CRS3123 for MetRS. (**D**) Ibezapolstat for polymerase IIIC. (**E**) Neither CRS3123 nor Ibezapolstat inhibit CamA at 10 μM. Error bars indicate the mean ± SD of N=3 independent determinations.

### SAM analogs as antibiotics

A known limitation of SAM analogs is their poor membrane permeability. In mammalian cells, SAM uptake may occur via nucleoside carriers ^120^, but SAM crosses cytoplasmic membranes poorly ^121^. In bacteria, SAM uptake appears to occur only via specific transporters, typically produced only by obligate parasites ^26, 122^. Prodrug strategies have been employed to improve cellular SAM uptake in human cells ^61, 73^, however it is unclear whether these strategies will be effective in penetrating bacterial cell walls.

Another consideration is possible development of resistance via suppressor mutations. A recent study reported that sinfungin exhibits antifungal activity by disrupting RNA adenine methylation, thereby impairing key pathogenic traits of *Candida albicans* – a fungus commonly found in the gastrointestinal tract and oral cavity – including hyphal morphogenesis, biofilm formation, and epithelial adhesion ^123^. However, an earlier study in the fungus *Saccharomyces cerevisiae* showed that resistance to sinefungin can arise through mutations in the high-affinity SAM transporter (Sam3) or through upregulation of SAM synthase and mRNA cap MTase ^124^. Possible bacterial resistance mechanisms to SAM analogs have not yet been well studied.

### Comparison to other inhibitors

Two orally-active small molecules have been developed and entered clinical trials for treating *C. difficile* infection: CRS3123, an inhibitor of bacterial methionyl-tRNA synthetase (MetRS) ^125, 126^ and ibezapolstat, which targets DNA polymerase IIIC in Gram-positive bacteria with low G+C content ^127, 128^. CRS3123 is a diaryldiamine compound featuring two bicyclic aromatic rings connected by a flexible diamine bridge (Figure 6C). The structure of CRS3123 bound to MetRS2 from the multidrug-resistant Gram-negative bacterium *Xanthomonas citri* has been solved ^129^ (PDB 6WQT) (Figure 6C). Ibezapolstat is a guanine analog with a morphilinoethyl group at the N7 position and a dichlorobenzylamine group at the C2 position (Figure 6D). Its proposed inhibitory mechanism involves competition with cognate dGTP at the active site of pol IIIC, pairing with the template cytosine; however, no structural data are currently available for ibezapolstat-bound DNA pol IIIC. Neither compound inhibits CamA at 10 μM (Figure 6E); nevertheless, we aim to incorporate chemical features - particularly from ibezapolstat (a guanine analog) - into the design of next generation adenosine analogs with enhanced cellular permeability.

In summary, our results provide chemical probes for exploring the role of CamA in sporulation and colonization, with potential as antivirulence agents against *C. difficile* infection. Our study also introduces the first chemical probes for inhibiting bacterial CcrM and human MettL5, each of which plays key roles in their respective hosts.

## METHODS

### Chemistry

Compounds **1**-**113** were synthesized by following our previously reported general methods ^73^. Sinefungin (**114**) was purchased from Sigma-Aldrich (Cat. No. S8559); CRS3123 (Cat. No. HY-18323) and ibezapolstat (Cat. No. HY-128357) were purchased from MedChemExpress. Compound MC4741 was reported previously ^64^.

### General procedure for synthesis of compounds 1 to 31

To a stirring solution of **1** or **9** (0.07 mmol), phenylboronic acid (0.09 mmol), dioxane (1.6 mL), and H_2_O (0.4 mL) were added to Pd(PPh_3_)_4_ (8.0 mg, 0.007 mmol) and K_2_CO_3_ (29 mg, 0.21 mmol). The resulting solution was stirred inside a microwave at 125 °C for 30 min. After the mixture was cooled, the volatiles were removed under vacuum. The residue was dissolved in H_2_O/methanol (1/1), filtered, and purified by HPLC (MeCN/H_2_O) to give **2−8** and **10−31** (45−75% yield).

### General procedure for synthesis of compound 32 to 44

To a stirring solution of **NH2-C2-Thioadenosine** (163 mg, 0.5 mmol) or **NH2-C3-Thioadenosine** (170 mg, 0.5 mmol) in 5 mL of anhydrous DMF, TEA (152 mg, 1.5 mmol) and isocyanate (0.6 mmol) were added at room temperature. After being stirred at room temperature for 1−4 h, the mixture was diluted with 100 mL of Et_2_O and filtered. Then, the residue was dissolved in 5 mL of methanol for purification with prepHPLC (MeCN/H_2_O) to afford **32−44** (56−88% yield).

### General procedure for synthesis of compound 45 to 67

Compound **45** and intermediate **benzylamine-C3-thioadenosine** were synthesized by following our previously reported methods ^73^. Compounds **46−67** (43−82% yield) were synthesized from **45** or **benzylamine-C3-thioadenosine** by following the same method as **2−31**.

### General procedure for synthesis of compound 68 to 78

Compounds **68−78** (31−64% yield) were synthesized from **2-chloro-6,7-dimethoxyquinazolin-4-amineC2-thioadenosine** by following the same method as **2−31**.

### General procedure for synthesis of compound 79 to 97

Compounds **79−97** (54−81% yield) were synthesized from **4-iodophenoxy-C2-thioadenosine** similar to **2−31**.

### General procedure for synthesis of compound 98 to 112

Compounds **98−112** (38−64% yield) were synthesized from **N6-phenylpropyl NH2-C3-thioadenosine** similar to **1** and **9**.

### Nucleic acid adenine MTases used in the study

We characterized and prepared the seven MTases used in the current study in our laboratories: *Escherichia coli* Dam (pXC1612) ^130, 131^, *Caulobacter crescentus* CcrM (pXC2121) ^83^, *Clostridioides difficile* CamA (pXC2184) ^77^, human MettL3–MettL14 ^114, 115^, human MettL16 (pXC2210) ^81^, human MettL5–Trm112 (pXC2062–pXC2076) ^81, 132^, and human PCIF1 (pXC2055) ^133, 134^. MettL3-MettL14 was expressed in Sf9 insect cells, and the other enzymes were expressed in *Escherichia coli* strain BL21(DE3).

### Inhibition assays

Methylation inhibition assays were conducted in the presence of 10 µM inhibitors, with detailed conditions summarized in Table 1. Assays were performed in low-volume 384-well plates containing 5 μl of reaction mixture per well, and luminescence was measured using a Synergy 4 multimode microplate reader (BioTek). Following the reactions, all samples were quenched by adding trifluoroacetic acid (TFA) to a final concentration of 0.1% (v/v). Methylation activity was assessed using the MTase-Glo bioluminescence assay (Promega) ^75^, which detects the reaction by-product SAH. SAH is enzymatically converted to ATP in a two-step reaction, and ATP levels are then quantified via a luciferase-based luminescence readout. For IC₅₀ determination, the same reaction setup was used as described in Table 1, with the exception that inhibitor concentrations were varied by two-fold series dilution.

### AlphaFold3 modeling of compounds binding in CcrM, MettL5-Trm112 and MettL3-MettL14

Protenix server (https://protenix-server.com/login) was utilized to generate five top hits of compounds 67, 37, and 77, respectively with CcrM, MetttL5-Trm112 and MettL3-MettL14. PyMol version 3.1.4.1 (Schrödinger, LLC) was used to prepare structure images.

## ASSOCIATED CONTENT

### Supporting Information

The supporting information contains molecular formula strings (SMILES) and associated inhibition data.

## AUTHOR INFORMATION

### Corresponding Authors

Rong Huang – Borch Department of Medicinal Chemistry and Molecular Pharmacology, Purdue Institute for Drug Discovery, Purdue Institute for Cancer Research, Purdue University, West Lafayette, Indiana 47907, United States;

Xiaodong Cheng − Department of Epigenetics and Molecular Carcinogenesis, University of Texas MD Anderson Cancer Center, Houston, Texas 77030, United States;

### Authors

Jujun Zhou − Department of Epigenetics and Molecular Carcinogenesis, University of Texas MD Anderson Cancer Center, Houston, Texas 77030, United States

Youchao Deng – Borch Department of Medicinal Chemistry and Molecular Pharmacology, Purdue Institute for Drug Discovery, Purdue Institute for Cancer Research, Purdue University, West Lafayette, Indiana 47907, United States

Dan Yu - Department of Epigenetics and Molecular Carcinogenesis, University of Texas MD Anderson Cancer Center, Houston, Texas 77030, United States

Xing Zhang − Department of Epigenetics and Molecular Carcinogenesis, University of Texas MD Anderson Cancer Center, Houston, Texas 77030, United States

Robert M. Blumenthal − Department of Medical Microbiology and Immunology, and Program in Bioinformatics, The University of Toledo College of Medicine and Life Sciences, Toledo, Ohio 43614, United States

### Authors Contributions

J.Z. performed inhibition assays, protein purifications of CamA, CcrM and PCIF1, and AF3-based structural modeling; Y.D. performed compound synthesis. D.Y. performed protein purifications of MettL5-Trm112 and MettL16. T.H and M.V. provided recombinant MettL3-MettL14 complex. R.M.B. assisted in preparing the manuscript and participated in discussions; X.Z. provided supervision, conceptualization and project administration; R.H. and X.C. designed and organized the study, performed writing, reviewing and editing of the manuscript, conceptualization, and funding acquisition.

## ACKNOWLEDGMENTS

The work was supported by U.S. National Institutes of Health grant R35GM134744 (to X.C., who is a CPRIT scholar in Cancer Research), and Purdue University Evanson-McCoy Endowment (to R.H.). We thank John R. Horton for providing Dam enzyme; Dr. Danti Rotili and Dr. Antonello Mai for providing compound MC4741.

## ABBREVIATIONS

CamA: *Clostridioides difficile-*specific DNA adenine methyltransferase
CDI: *Clostridioides difficile* infections
IC_50_: half maximal inhibitory concentration
*K*_D_: dissociation constant
MTase: methyltransferase
PRMT: protein arginine methyltransferase
SAH: *S*-adenosyl-*L*-homocysteine
SAM: *S*-adenosyl-L-methionine
TFA: trifluoroacetic acid

